# Performance analysis of novel toxin-antidote CRISPR gene drive systems

**DOI:** 10.1101/628362

**Authors:** Jackson Champer, Isabel Kim, Samuel E. Champer, Andrew G. Clark, Philipp W. Messer

**Author notes:** Corresponding author: JC.

## Abstract

Gene drives can potentially fixate in a population by biasing inheritance in their favor, opening up a variety of potential applications in areas such as disease-vector control and conservation. CRISPR homing gene drives have shown much promise for providing an effective drive mechanism, but they typically suffer from the rapid formation of resistance alleles. Even if the problem of resistance can be overcome, the utility of such drives would still be limited by their tendency to spread into all areas of a population. To provide additional options for gene drive applications that are substantially less prone to the formation of resistance alleles and could potentially remain confined to a target area, we developed several designs for CRISPR-based gene drives utilizing toxin-antidote (TA) principles. These drives target and disrupt an essential gene with the drive providing rescue. Here, we assess the performance of several types of TA gene drive systems using modeling and individual-based simulations. We show that Toxin-Antidote Recessive Embryo (TARE) drive should allow for the design of robust, regionally confined, population modification strategies with high flexibility in choosing drive promoters and recessive lethal targets. Toxin-Antidote Dominant Embryo (TADE) drive requires a haplolethal target gene and a germline-restricted promoter but should enable the design of both faster regional population modification drives and even regionally-confined population suppression drives. Toxin-antidote dominant sperm (TADS) drive can be used for population modification or suppression. It spreads nearly as quickly as a homing drive and can flexibly use a variety of promoters, but unlike the other TA systems, it is not regionally confined and requires highly specific target genes. Overall, our results suggest that CRISPR-based TA gene drives provide promising candidates for further development in a variety of organisms and may allow for flexible ecological engineering strategies.

## INTRODUCTION

A successful gene drive will rapidly spread through a population by biasing inheritance in favor of the drive allele^1–7^. This can be used for population modification, or with an appropriate drive arrangement, population suppression^1–7^. The potential applications of such drives are numerous, with perhaps the most promising involving modification or suppression of mosquito populations to prevent transmission of vector-borne diseases such as malaria or dengue^1–3,5^. Similar techniques can potentially be used against invasive species or agricultural pests^1–3,5^.

Homing gene drive constructs based on CRISPR-Cas9 have been constructed in yeast^8–11^, flies^12–18^, mosquitoes^19–21^, and mice^22^. These constructs work by cleaving a wild-type allele, resulting in copying of the drive allele during homology-directed repair. If cleavage is repaired by end-joining, however, mutations can be created at the guide RNA (gRNA) target site^15^. This often results in the formation of a “resistance allele”, since the mutated target site prevents future cleavage by the homing drive and thus impairs its spread. Such resistance alleles can form in the germline as an alternative to homology-directed repair or during early embryo development by maternally-deposited Cas9 and gRNA^15^. Some strategies for reducing resistance allele formation have already been successfully tested, including gRNA multiplexing^16^, improved promoters^16,23^, and selection of a highly conserved target site where mutations are not tolerated^24^. This latter method, when combined with an improved promoter, recently resulted in the successful suppression of *Anopheles gambiae* in population cages.

Though promising, such a strategy may be difficult to apply to population modification gene drives. This is because it relies on changed resistance allele sequences rendering the target gene non-functional, allowing them to contribute to the overall goal of population suppression, even if they slow the spread of the drive. A population modification drive would only be able to remain viable while removing resistance alleles if it targets an essential gene and itself has a recoded version of the target that is fully functional. This requires targeting a site that can be recoded, which allows the possibility of forming resistance alleles that do not disrupt the function of the target gene. A homing drive with multiplexed gRNAs all targeting within a narrow window could likely overcome this limitation while retaining high drive efficiency, but such drives may still be vulnerable to formation of resistance alleles by partial homology-directed repair that copies the recoded region but not the payload gene, essentially forming a functional resistance allele. Aside from this limitation, all homing drives also require Cas9 cleavage specifically in the germline to allow for homology-directed repair instead of end-joining, which otherwise predominates, particularly when cleavage occurs in the embryo. This requires choosing a suitable promoter, which may often be difficult to find in new species and could thereby prove to be a barrier to development of homing drives.

Another inherent feature of homing gene drives that could limit their utility is that they are rapidly spreading constructs that cannot be easily confined to a specific geographic region^25^. In fact, the migration of only a few individuals could be sufficient to establish a homing drive in a new population, which could be particularly undesirable when the goal is suppression of invasive species or agricultural pests outside their native range^26^. Thus, new gene drive options are needed that are effective, flexible, and can be regionally confined.

One possible strategy for developing an efficient drive is to avoid the need for homology-directed repair after cleavage by using a toxin-antidote (TA) drive system. This principle is often seen in natural drives^27^ and has been successfully applied for the *Medea* system^28^. However, *Medea* elements proved to be highly specific to *Drosophila* and difficult to transfer into other species. CRISPR nucleases could in principle be used to create highly flexible systems, where the “toxin” would consist of Cas9 and gRNAs targeting an essential gene. The “antidote” would then be a recoded copy of the gene immune to drive cleavage. With both the toxin and antidote as part of the drive allele, it would steadily convert wild-type alleles to disrupted alleles in the population, which would systematically be removed from the population, thereby increasing the relative frequency of the drive over time.

There are several potential variants of TA systems, including those based on recessive lethal genes (TARE, Toxin-Antidote Recessive Embryo), haplolethal genes (TADE, Toxin-Antidote Dominant Embryo), and genes with expression after meiosis I that is required for sperm development (TADS, Toxin-Antidote Dominant Sperm). Recently, we developed a “same-site” TARE drive capable of spreading through population cages^29^, and others constructed a “distant-site” TARE drive termed ClvR that was similarly successful^30^. Thus, such systems are highly promising for further development. Here, we analyze the expected population dynamics of TARE, TADE, TADS, and several variants of these strategies using modeling and simulations. In different forms, such systems are capable of providing for highly efficient population modification or suppression strategies, either of which could be global or regional, depending on the specific system.

## METHODS

### Deterministic model

To analyze the dynamics of TA systems, we developed a deterministic, discrete-generation modeling approach. These models were used to demonstrate frequency trajectories and to calculate parameters such as introduction thresholds. Initially, drive/wild-type heterozygotes are added to a population of wild-type individuals at a specified introduction frequency. In each generation, frequencies were tracked for each genotype. Females select a mate randomly, with each male’s chance of being selected being proportional to his fitness value. Females then generate a number of potential offspring equal to twice their fitness value.

Individuals homozygous for a drive allele (or with a drive allele on the Y chromosome) carry a fitness cost that reduces fecundity for females and the probability of being selected as a mate for males. Drive heterozygotes are assumed to have a fitness equal to the square root of the fitness of homozygous individuals (i.e. we assume that fitness costs of the drive allele are multiplicative). Several events can take place in the model depending on the particular drive strategy, including successful germline drive conversion for homing drive heterozygotes, removal of non-viable sperm in males due to a TADS system or X-shredder, and successful disruption of the target gene in the germline and embryo (such disruption does not take place for homing drives due to a drive-optimized promoter, or in the embryo for TADE and X-shredder drives). Offspring with non-viable genotypes due to toxin effects are then removed. Genotype frequencies are finally renormalized to produce the population state for the following generation.

### Gene drive simulations

To gain a better understanding of how our TA gene drive designs would perform in more realistic models, we developed individual-based simulations that incorporated a degree of stochastic variation to model the effects of genetic drift. These simulations were used to determine average drive performance based on different initial parameters (all heat maps shown in this study are based on these simulations).

All simulations were implemented in the forward-in-time population genetic simulation framework SLiM version 3.2.1^31^. Our basic model simulates a panmictic population with males and females with discrete, non-overlapping generations. To obtain the individuals of the next generation, each female randomly selects a mate, with drive males having a reduced probability of being selected as in the deterministic model if the drive allele has a fitness cost. We then calculate the number of offspring generated based on a binomial distribution with maximum of 50 and *p* = fitness/25, so that a female with fitness = 1 will have an average of two offspring. Fitness is determined by genotype as described in the deterministic model and is multiplied by a density dependent factor equal to 10/(1+9*N*/*K*) where *N* is the total population and *K* is the carrying capacity. This density factor was selected to produce logistic dynamics and to smoothly but quickly restore the population to carrying capacity after perturbation, unless a population suppression system produces downward pressure on the population. At low densities, it produces a maximum 10-fold population growth rate per generation, thus allowing for rapid growth when individuals are not limited by competition.

The next step is to generate offspring. Each offspring randomly receives an allele from each parent. If the target allele was wild-type and the parent also had a drive allele, then the wild-type allele is converted to a disrupted allele with a probability equal to the germline cut rate. If the offspring has any wild-type alleles after parental germline cutting and has a mother with at least one drive allele, then any wild-type alleles are converted to disrupted alleles with a probability specified by the embryo cut rate. For the TADS drive, if the offspring received a disrupted allele from the male parent, the genotype of the child is redrawn, since such an allele would be incapable of fertilizing an egg. Finally, offspring with non-viable genotypes are removed from the population.

In our simulations, we assumed that drive individuals were first mated with wild-type individuals to produce heterozygous offspring, which were then introduced at a frequency representing 20% of the total population (for TADS Y-linked suppression drive, all introduced individuals were males with the drive). Several drive performance parameters were fixed at standardized levels inspired by laboratory gene drive mosquitoes^20,24^, unless otherwise specified. These include 99% germline cut rate, 95% embryo cut rate (5% for TADE and TADE suppression drives, which are intolerant of high embryo cut rates - such low embryo cut rates have also been achieved in gene drive mosquitoes^23,24^), and 95% drive homozygote fitness compared to wild-type individuals. These parameters are then varied in our analyses (individually or in combination) to study how they affect drive dynamics. Each simulation had a starting population of 100,000 individuals and was evaluated over 100 generations.

### Data generation

Simulations were run on the computing cluster of the Department of Computational Biology at Cornell University. Data processing, analyses, and figure preparation of simulations were performed in Python and R. All SLiM configuration files for the implementation of the simulations and all data are available on GitHub (https://github.com/MesserLab/ToxinAntidoteSystems). All simulations were replicated a total of ten times for each parameter setting, and the results were averaged.

## RESULTS

### General TA drive principles

TA drive alleles each contain a toxin and an antidote that will rescue the effect of the toxin. We assume that the toxin is a CRISPR nuclease targeting an essential gene that will get disrupted when the break is repaired by end-joining or homology-directed repair through introduced mutations at the cut sites, rendering it non-functional. The antidote consists of a recoded version of the gene that no longer matches the gRNAs and therefore, cannot be disrupted by the drive. Cells or individuals exposed to the toxin will thus perish, unless rescued by a drive allele (Figure 1A), with details determined by the specific type of drive. Various potential arrangements and targets can be conceived, depending on whether the goal is population modification (Figure 1B) or population suppression (Figure 1C), which we will describe in the individual sections below. TARE and Toxin-Antidote Double Dominant Embryo (TADDE) drive systems are expected to reach all individuals quickly in idealized population models (Figure S1). However, it can take an extended time for these constructs to reach their maximum value (Figure 1B), dependent on the drive’s fitness cost (Figure 1D). Additionally, most TA systems have introduction frequency thresholds below which the drive will not spread at all (Figure 1D), except for the TADS drive, which should have a zero-threshold and rapidly spread through populations from even a small initial frequency.

**Figure 1.**
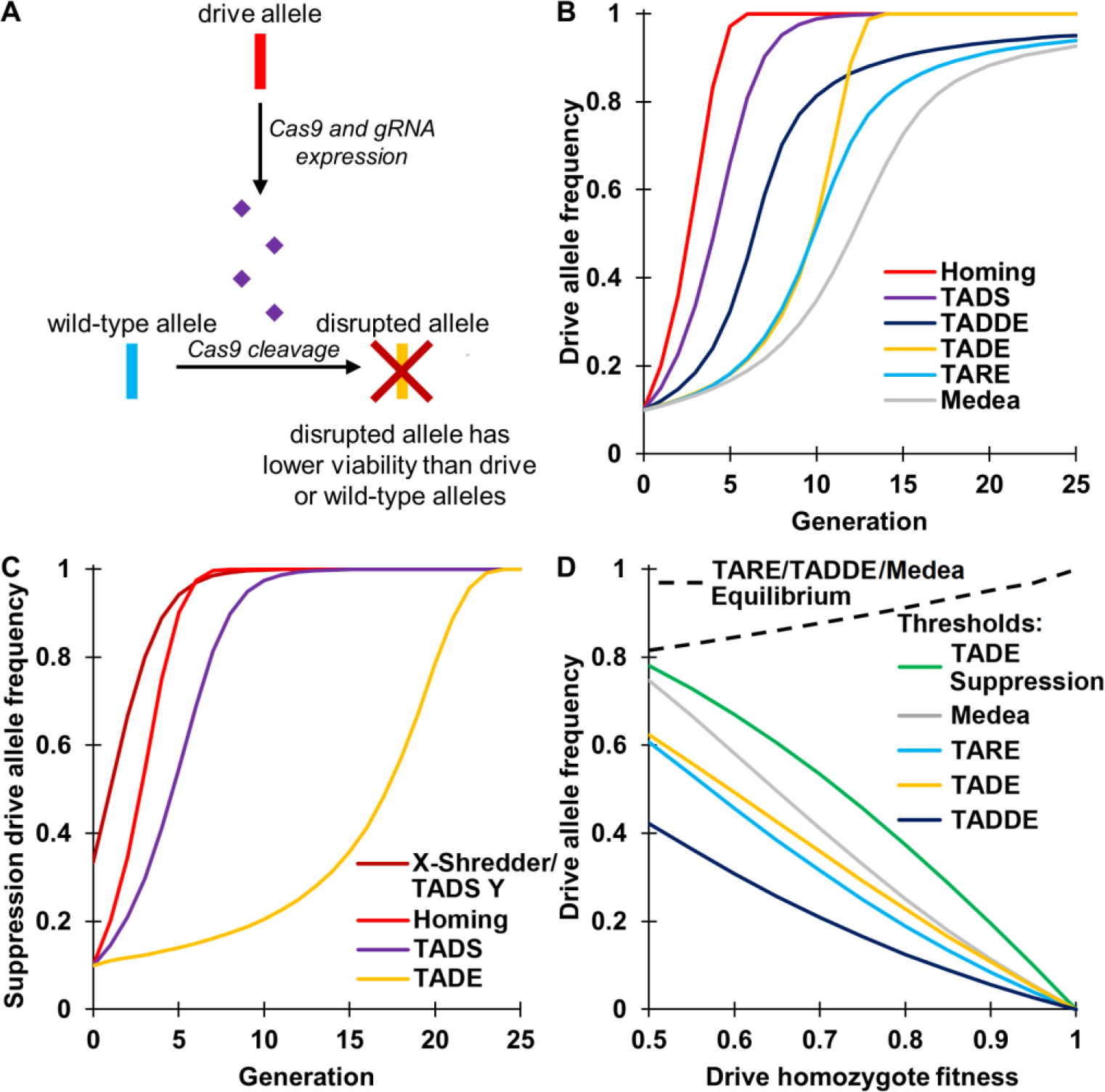
Overview of TA systems and comparison to other drives. (**A**) The general principle behind TA systems is disruption of wild-type alleles by the drive, which are eventually rendered non-viable. The drive remains viable, increasing its relative frequency in the population. (**B**) Expected allele frequencies when drive heterozygotes are released into a wild-type population at 20% starting frequency (10% starting drive allele frequency) in a discrete-generation, deterministic model. All drives are assumed to have no fitness cost and 100% drive efficiency, with no resistance formation. (**C**) In some forms, TA drives can induce population suppression (model as in **B**). (**D**) TARE, TADE, TADDE, and *Medea* drives have fitness-dependent introduction thresholds, above which the drive will increase in frequency and below which it will decrease. TARE, TADDE, and *Medea* drives additionally have final equilibrium frequencies that are dependent on fitness costs. At equilibrium, all individuals carry at least one copy of the drive, but some carry disrupted alleles as well.

### TARE drive

TARE drives target a haplosufficient but recessive lethal gene, with the drive providing rescue (Figure 2A). It is thus desirable that nuclease cleavage occurs not only in the germline, but also in the early embryo due to maternally deposited Cas9 and gRNA. With a highly efficient drive, female heterozygotes crossed with wild-type males will therefore have only drive-carrying offspring. This is because offspring that don’t inherit a drive allele will inherit a disrupted target allele from the mother and a wild-type allele from the father, which will then typically become disrupted due to maternal Cas9 activity, rendering the individual non-viable. Because this drive does not work by increasing the number of drive alleles, but by reducing the number of wild-type alleles (thus increasing the relative frequency of the drive), it shows threshold-dependent dynamics (Figure 1B, Figure 1D, Figure 2B). This drive should be highly tolerant of variation in expression from the nuclease promoter, with optimal performance when both germline and embryo cut rates are high (Figure 2C). Indeed, the promoter need not even be restricted to driving expression in the germline and early embryo. Constitutively active promoters would presumably work equally well (though they may have a slightly higher fitness cost), as long as there is expression in germline or germline precursor cells.

The TARE drive can be “same-site” as in Figure 2A or a “distant-site” drive in which the drive allele is not located at the same genomic site as the target allele (Figure S2A). Successful same-site^29^, and distant-site^30^ systems have already been engineered with high germline and embryo cut rates and little to no observable fitness costs. These systems have nearly equivalent performance when cut rates are high (Figure S2B), but the distant-site drive retains higher performance when both the germline and embryo cut rates are low (Figure S3C) since it often has two wild-type alleles available to cleave rather than just one as for the same-site drive. On the other hand, a same-site drive may be easier to engineer since the recoded region may be smaller and the natural target gene promoter would still drive expression of the rescue allele. The natural promoter and genomic site of the rescue element may also avoid the pitfall of incomplete rescue that would be a more significant consideration for distant-site drives.

In our model, TARE systems reach all individuals quickly with a modest release size (Figure S1), but their rate of increase becomes slowed at high frequencies (Figure 1B), which could be an issue for a population modification strategy where the payload is substantially more effective in homozygotes than heterozygotes. To avoid this, the target of a TARE system could be located on the X-chromosome, which would make males with only one copy of the disrupted target gene non-viable (Figure S3D). This would allow the drive allele to fixate substantially more quickly than autosomal TARE systems (Figure S3E). However, X-linked TARE drives would not have any cleavage activity in the germline of males and thus, have a slower rate of spread than autosomal systems (Figure S3F), at least until the drive reaches all individuals.

**Figure 2.**
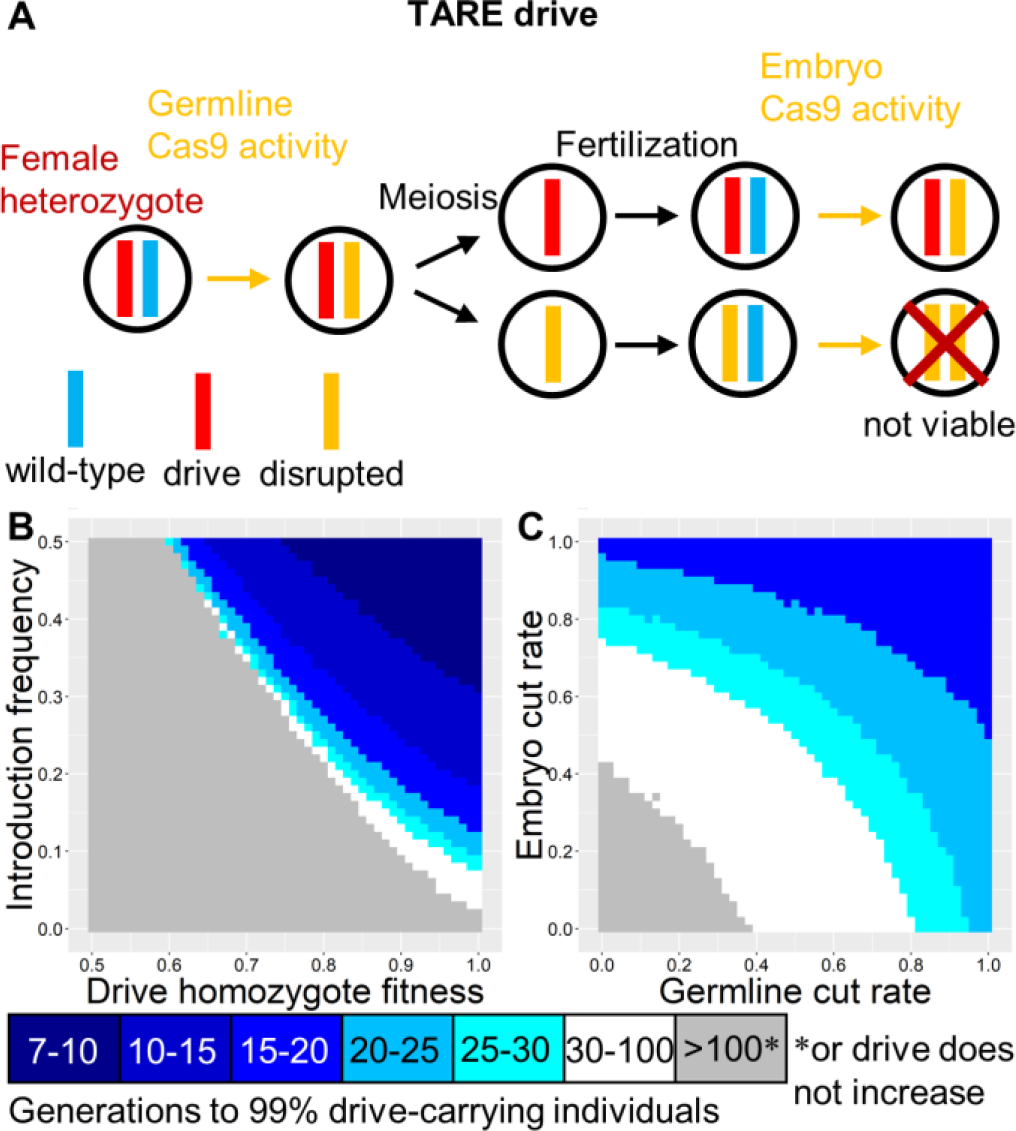
TARE drive. (**A**) In the TARE drive, germline activity disrupts the target gene, followed by embryo activity in the progeny of drive-carrying females. The target gene is recessive lethal, so any individuals inheriting two disrupted target genes are non-viable. By contrast, all individuals with a wild-type or drive allele are viable. (**B**) The speed at which the TARE drive reaches 99% of individuals in the population with varying introduction frequency and drive fitness. (**C**) Same as **B**, but with varying germline and embryo cleavage rates. Grey means that the drive failed to reach 99% because it spread too slowly or was not able to spread at all.

### TADE drive

TADE drives target a haplolethal gene, with the drive providing rescue (Figure 3A). Like TARE, such a drive is expected to show threshold-dependent dynamics (Figure 1B, Figure 1D, Figure 3B). For a TADE drive, nuclease cleavage should occur only in germline gametocytes. Otherwise drive/wild-type heterozygotes will not have two functioning copies of the haplolethal gene in all cells, which will likely cause death, depending on the magnitude of expression outside the germline. Similarly, embryo cleavage activity would render drive-carrying individuals non-viable. Though the promoter is germline-restricted, it could still have expression before or after the narrow window in early meiosis for homology-directed repair, allowing somewhat more tolerance in the specificity of expression timing than for promoters used in homing drives. With a suitable promoter, the offspring of both males and females that fail to inherit the drive will perish. This allows the TADE drive to spread more rapidly than the TARE drive and quickly reach fixation (Figure 1B). However, substantial embryo resistance would likely prevent the drive from spreading (Figure 3C).

As with the TARE drive, the TADE drive can be “same-site” or “distant-site” (Figure S3A). These are expected to have similar performance (Figure S3B), but the distant-site drive may remain viable for higher embryo resistance rates when germline cleavage is low (Figure S3C). This is because a low rate of embryo cleavage events can help remove wild-type alleles that were not cleaved in the germline due to low germline cleavage rates. The drive alleles in this situation should still remain viable in most instances, since the other wild-type target allele should often remain undisrupted.

**Figure 3.**
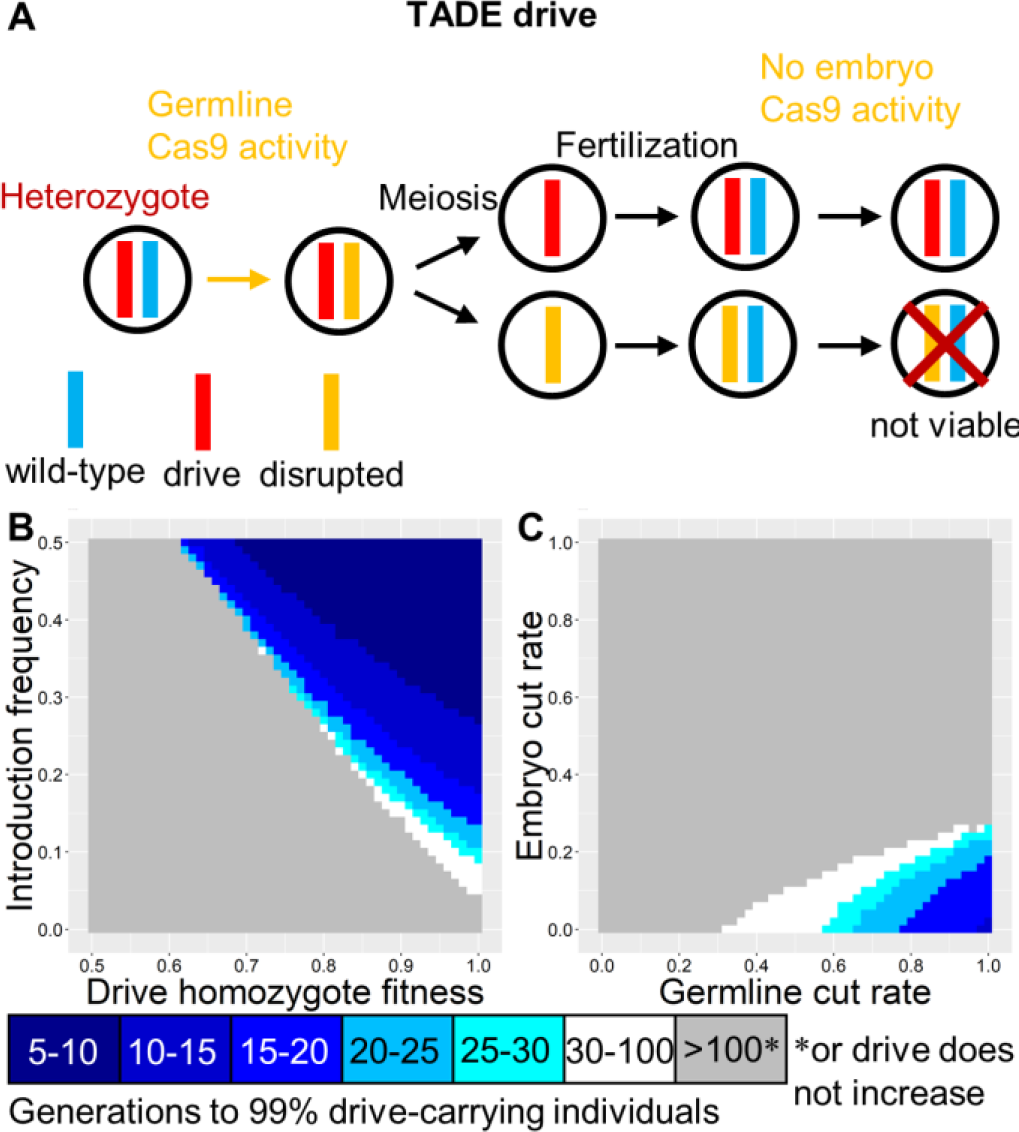
TADE drive. (**A**) In the TADE drive, germline activity disrupts the target gene, and the nuclease promoter is selected to minimize embryo activity. The target gene is haplolethal, so any individuals inheriting one disrupted target allele will be non-viable, even if the other allele is drive or wild-type. (**B**) The speed at which TADE drives reach 99% of individuals in the population with varying introduction frequency and drive fitness. (**C**) Same as **B**, but with varying germline and embryo cleavage rate.

### TADE suppression drive

The TADE suppression drive is a form of distant-site TADE in which the drive is located in a recessive female (or male, but not both) fertility (or viability) gene, disrupting the gene with its presence (Figure 4A). Thus, female drive homozygotes are sterile. If the germline cleavage rate is less than 100%, this drive would not fixate but instead impose a suppression factor on the population (Figure 4B), which is the fold-reduction in the population between generations after the drive reaches it maximum frequency, assuming no fitness effects on individuals due to reduced competition at low densities. Complete suppression occurs when the suppression factor is greater than the maximum population growth rate experienced by individuals at lower densities with less competition, before Allee effects dominate^32^. For the germline cut rates that have been observed experimentally in mosquito and *Drosophila* systems^12–21^, this would likely be sufficient to cause complete suppression. High suppression factors are also possible even if the target gene shows only partial haploinsufficiency (Figure S4). Note that unlike homing-based drives and X-shredders, the TADE suppression system shows threshold-dependent dynamics (Figure 1C, Figure 1D, Figure 4C), making it a regionally confined system. In an effective TADE suppression drive, the parameter space for embryo and germline cut rates is even more restricted than for TADE drive (Figure 4D), though still within the range demonstrated in mosquito drives.

**Figure 4.**
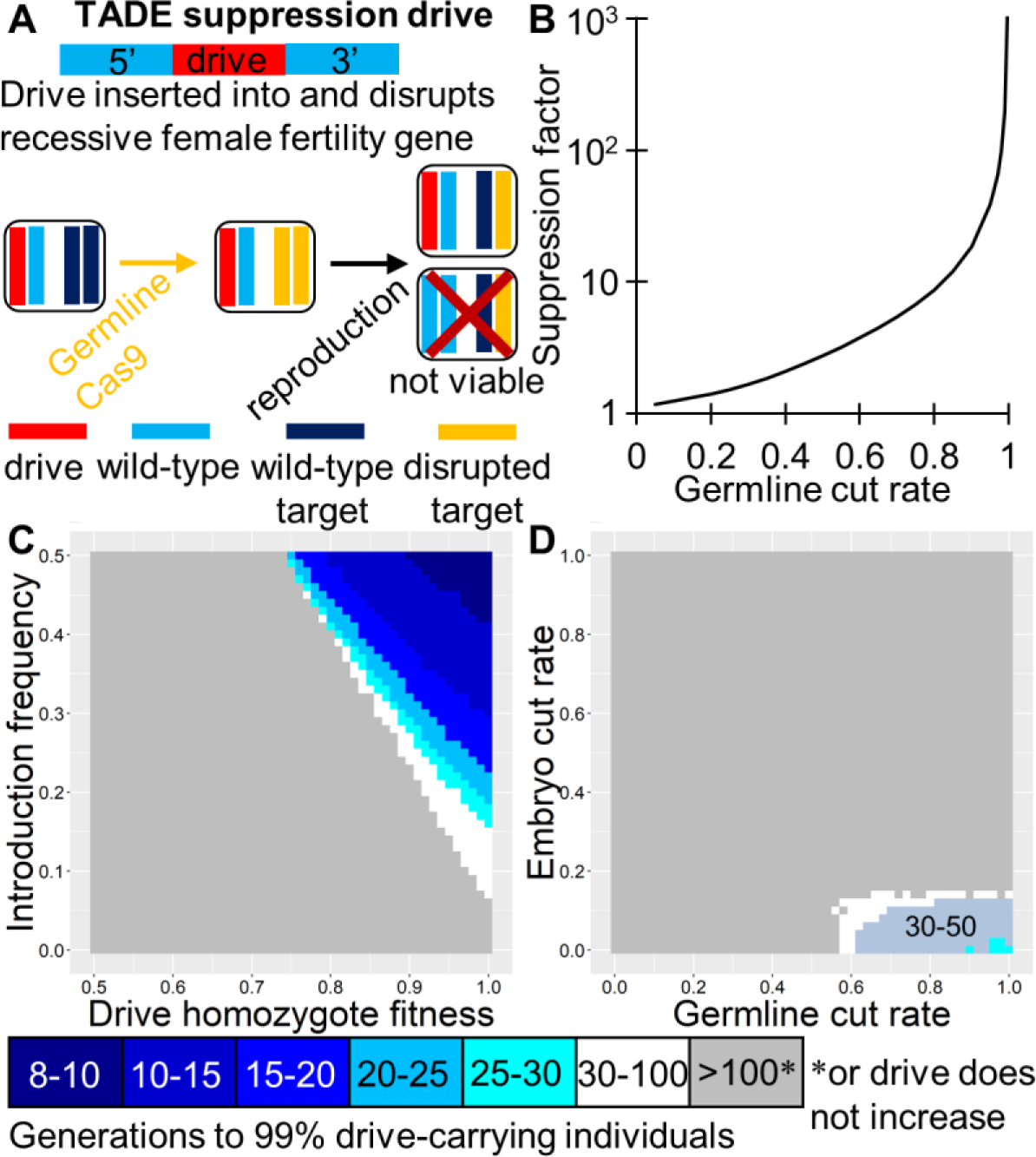
TADE suppression drive. (**A**) The TADE suppression drive is “distant-site’, so its target gene is unlinked from the drive allele, which is located in a female (or male) fertility gene. The drive disrupts the fertility gene, so female drive homozygotes are sterile. Germline activity disrupts the target gene, and the nuclease promoter is selected to minimize embryo activity. The target gene is haplolethal, so any individuals inheriting less than two wild-type target alleles and/or drive alleles are non-viable. (**B**) The ability of the TADE suppression drive to suppress a population. If the germline cleavage rate is 100%, suppression will occur. Otherwise, suppression will only occur if the suppression factor is greater than the fitness advantage of individuals at low population density. (**C**) The speed at which the TADE suppression drive reaches 99% of individuals in the population with varying introduction frequency and drive fitness. (**D**) Same as **C**, but with varying germline and embryo cleavage rate.

### TADDE drive

TADDE drives are simply TADE drives in which the rescue element either has two recoded copies of the haplolethal gene or a sufficiently altered promoter to increase expression of the rescue element. Thus, a single drive allele is sufficient to provide rescue even if paired with a disrupted allele (Figure 5A). TADDE thus allows for removal of wild-type alleles immediately after disruption in both males and females, while preventing removal of drive-carrying individuals, which occurs in TADE drive offspring when two drive heterozygotes mate. This should allow it to spread more quickly (Figure 2B) with a lower threshold (Figure 1D) than TADE or TARE systems, while otherwise still retaining similar dynamics (Figure 5B). Because drive alleles are not removed when paired with disrupted targets, embryo resistance can be fully tolerated (as well as somatic expression, as in TARE), though embryo cleavage does not significantly increase the rate of spread of this drive when germline cleavage is high (Figure 5C). Same-site and distant-site configurations of TADDE drive should be very similar except when both germline and embryo cleavage rates are exceptionally low (Figure S5).

**Figure 5.**
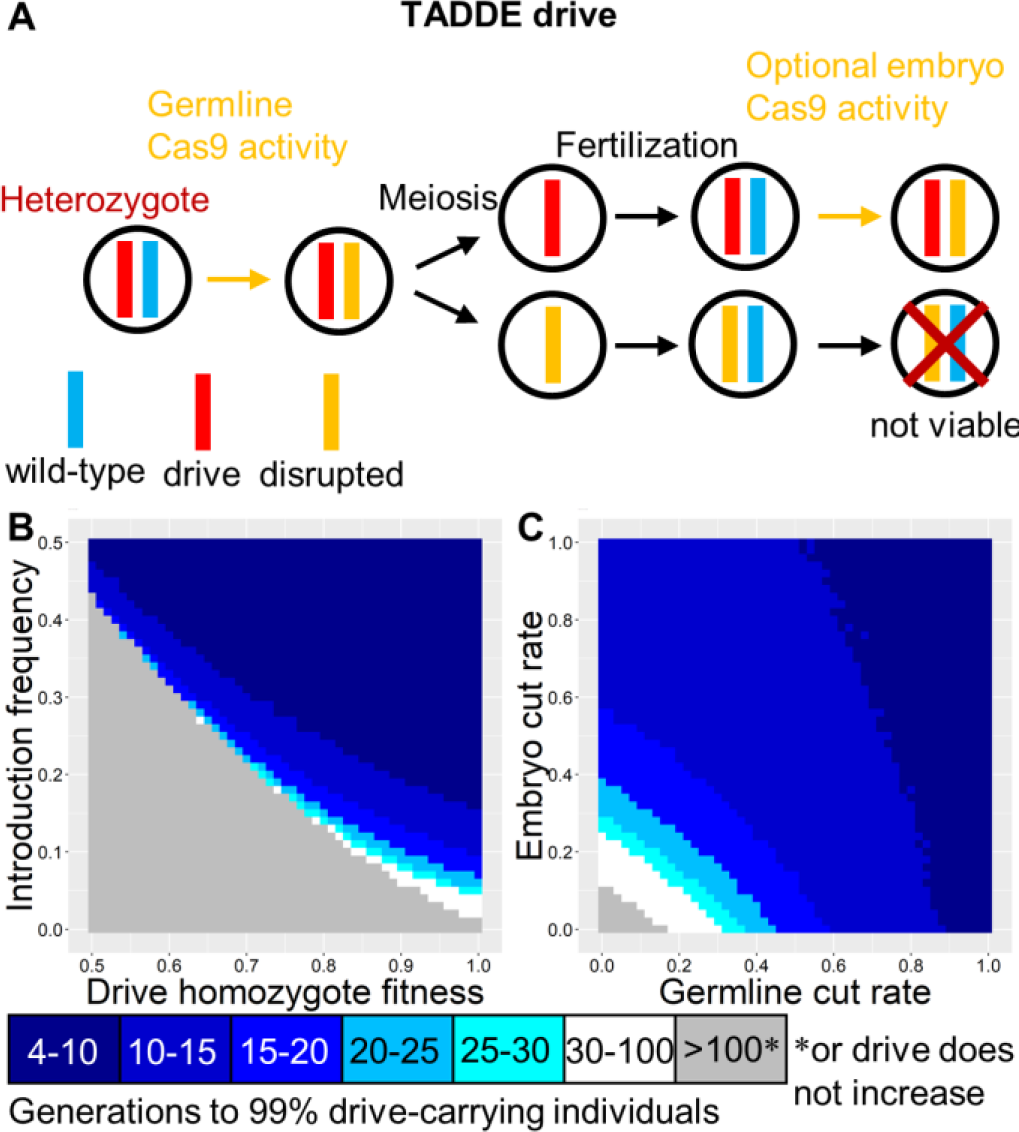
TADDE drive. (**A**) In the TADDE drive, germline activity disrupts the target gene, and embryo activity is optional (and preferred unless germline cleavage is 100%). The target gene is haplolethal, so any individuals inheriting at least one disrupted target allele are non-viable unless they also inherit a drive allele, which encodes two copies of the gene or provides sufficient expression of the target gene such that only one copy is needed for rescue. (**B**) The speed at which the TADDE drive reaches 99% of individuals in the population with varying introduction frequency and drive fitness. (**C**) Same as **B**, but with varying germline and embryo cleavage rate.

### TADS drive

TADS drives target a gene that is expressed in gametocytes after meiosis I in males, and this expression must be critical for successful spermatogenesis such that sperm with a disrupted target allele are non-viable. Thus, only sperm with drive or wild-type alleles will successfully fertilize eggs (Figure 6A). This is expected to directly increases the frequency of the drive allele in the population, allowing for rapid spread of the drive (Figure 1B). However, the rate of spread would be somewhat reduced if females can mate with multiple males and sperm compete to fertilize eggs. Because the frequency of the drive allele directly increases, the drive has a zero-introduction threshold (Figure 6B) and would therefore not be expected to remain regionally confined. Somatic expression would likely be fully tolerated for such a drive, and it should also allow for wide variety of promoters varying in both germline and embryo cut rates (Figure 6C), yet finding a suitable target gene will likely be difficult. In distant-site configuration (Figure S6A), TADS drive is similar to the same-site drive, although as with the other types of drive, distant-site TADS should perform better than same-site TADS when both germline and embryo cleavage rates are very low (Figure S6B-C).

**Figure 6.**
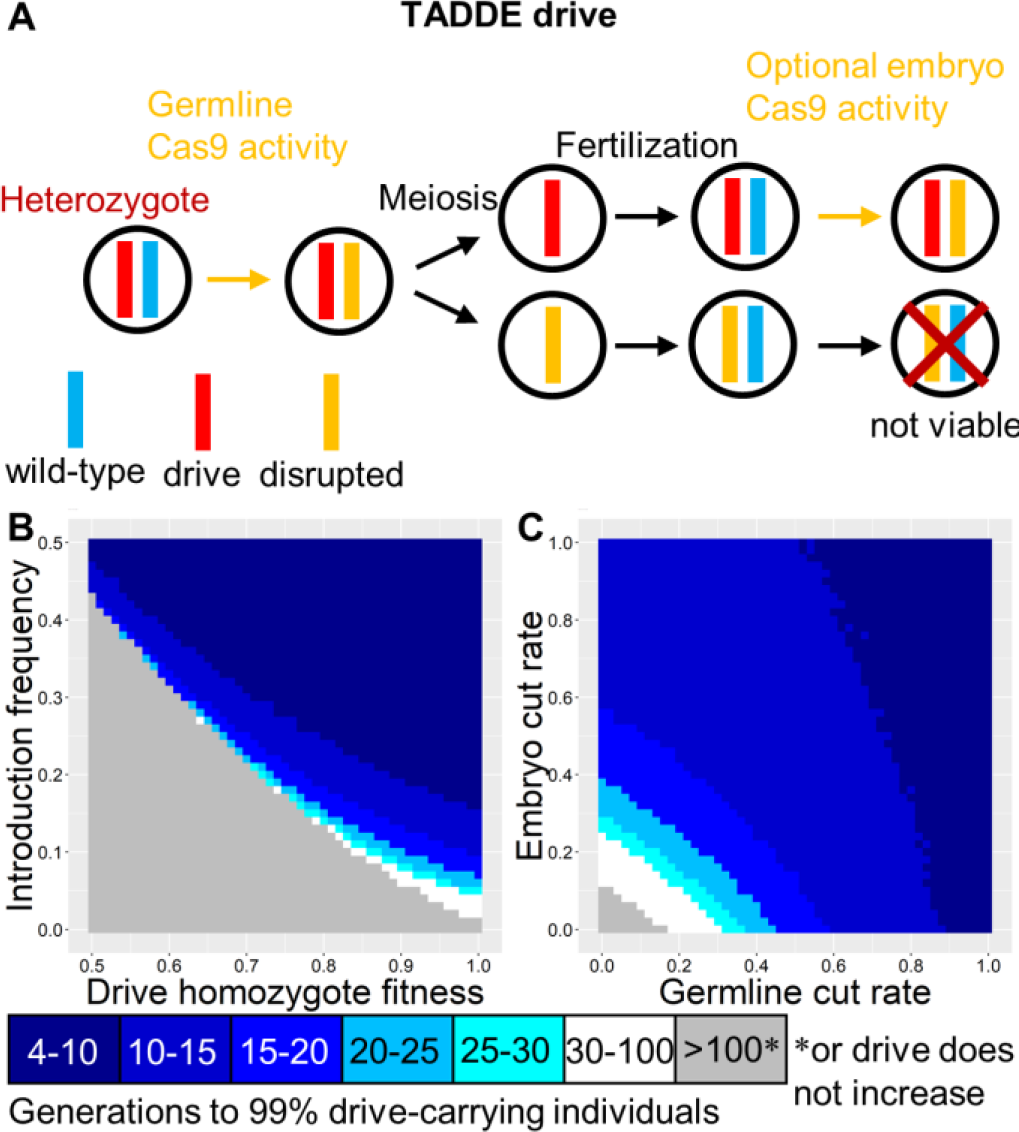
TADS drive. (**A**) In the TADS drive, germline activity disrupts the target gene, followed by embryo activity in the progeny of drive-carrying females. The target gene has expression in male gametocytes after meiosis I, and such expression is necessary for development of a viable sperm. Thus, sperm with disrupted alleles are non-viable. All sperm with a wild-type or drive allele are viable. (**B**) The speed at which the TADS drive reaches 99% of individuals in the population with varying introduction frequency and drive fitness. (**C**) Same as **B**, but with varying germline and embryo cleavage rate.

### TADS suppression drive

A distant-site TADS drive can be configured for population suppression by placement in a recessive male fertility (or viability) gene, disrupting the gene with its presence (Figure 7A). Thus, male drive homozygotes would be sterile. Because the drive works during spermatogenesis, it would be unable to provide any substantial suppression if located in a female (or both-sex) fertility or viability gene. However, in a male fertility gene, it would be expected to cause complete population suppression with zero introduction threshold (Figure 7B) nearly as rapidly as homing drives and X-shredders (Figure 1C). The suppression form of TADS is less tolerant of low embryo and germline cut rates than modification TADS systems, but it can still achieve success over a wide range of values (Figure 7C).

**Figure 7.**
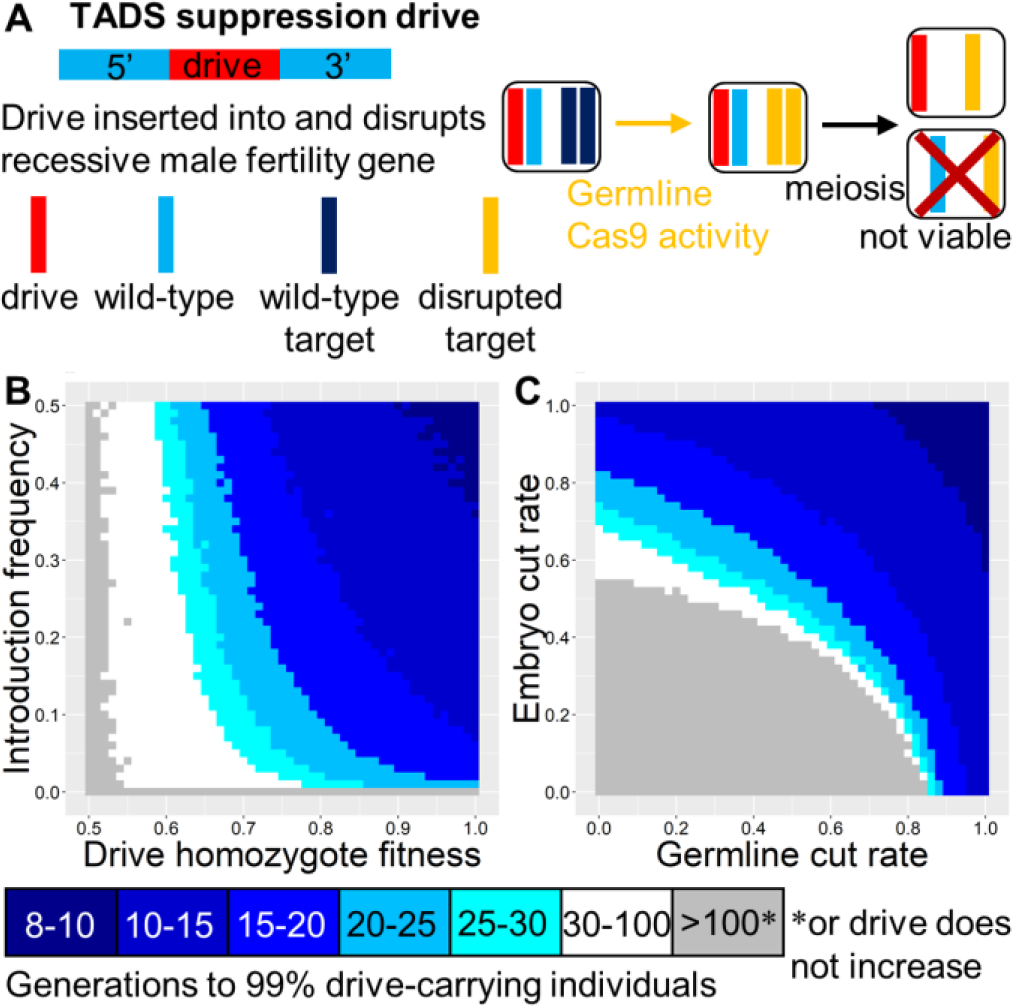
TADS suppression drive. (**A**) The TADS suppression drive is “distant-site’, located in a male fertility gene and with a target gene that is unlinked from the drive allele. The drive disrupts the fertility gene, so male drive homozygotes are sterile. Germline activity disrupts the target gene, followed by embryo activity in the progeny of drive-carrying females. The target gene has expression in male gametocytes after meiosis I, and such expression is necessary for development of a viable sperm. Thus, sperm with a disrupted target allele are non-viable unless they also have a drive allele. (**B**) The speed at which the TADS suppression drive reaches 99% of individuals in the population with varying introduction frequency and drive fitness. Full suppression would occur within a few generations of this point. (**C**) Same as **B**, but with varying germline and embryo cleavage rate.

### TADS Y-linked suppression drive

If a distant-site TADS drive is located on the Y chromosome (with an autosomal target), it will bias inheritance in favor of males (Figure 8A). This is expected to induce a germline cut rate-dependent suppression factor on a population (Figure 8B) after the drive fixates. According to our deterministic models, the suppression factor should be twice that of an X-shredder with a similar X-shredding rate. The dynamics of an ideal TADS Y-linked suppression drive should otherwise be almost identical to that of an ideal X-shredder (Figure 1C). The drive has a zero-threshold introduction frequency and is highly tolerant of both fitness costs (Figure 9C) and low germline cut rates (Figure 9D), though the germline cut rate will still need to induce a sufficient suppression factor if complete suppression is desired. A TADS suppression system could also be located on the X-chromosome, similarly biasing inheritance in favor of females and eventually inducing population suppression.

**Figure 8.**
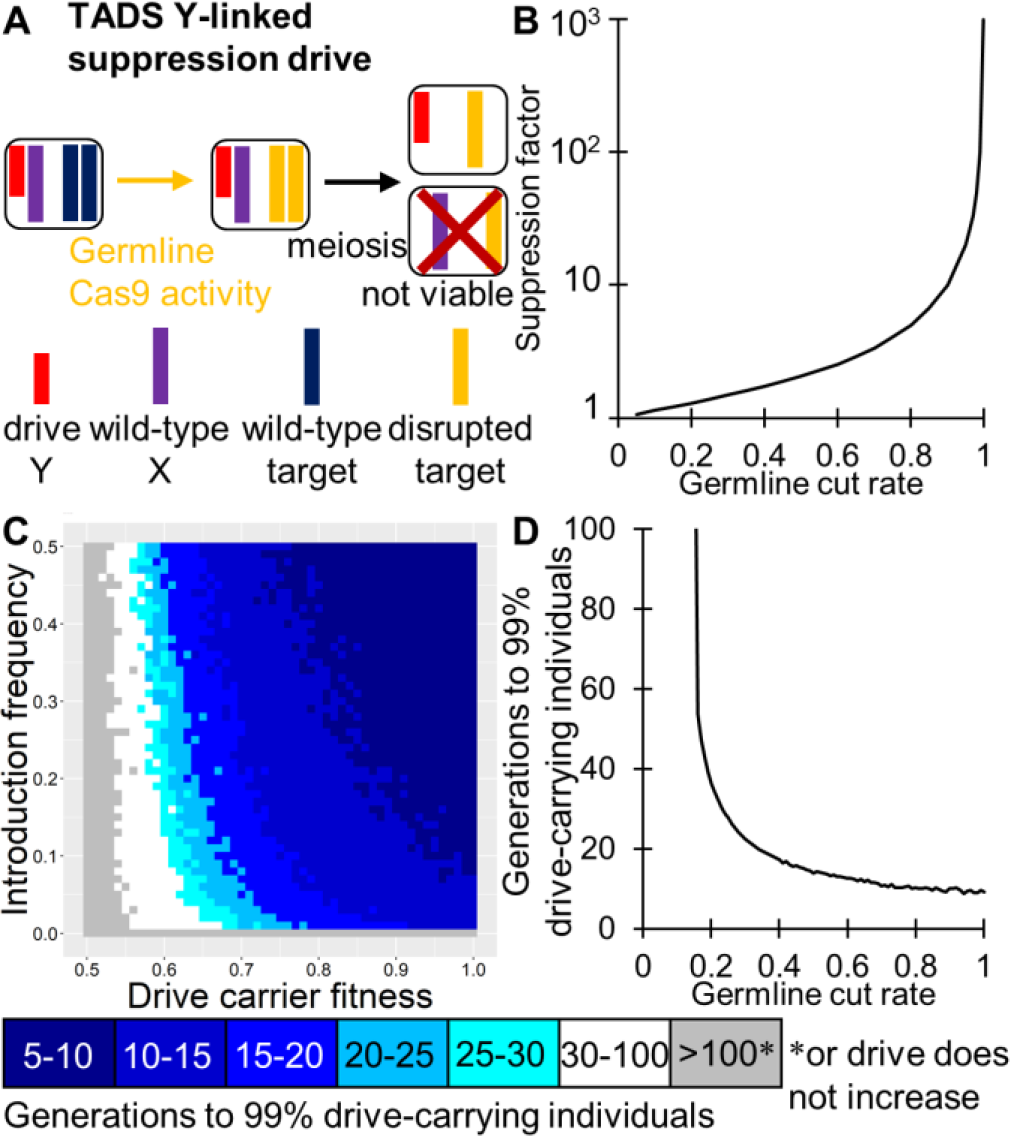
TADS Y-linked suppression drive. (**A**) The TADS Y-linked suppression drive is “distant-site’. It is located on the Y-chromosome and has a target gene that is not linked to the drive allele. Germline activity disrupts the target gene, followed by embryo activity in the progeny of drive-carrying females. The target gene has expression in male gametocytes after meiosis I, and such expression is necessary for development of a viable sperm. Thus, sperm with a disrupted target allele are non-viable unless they also have a drive allele. (**B**) The ability of the TADS Y-linked suppression drive to suppress a population. If the germline cleavage rate is 100%, suppression will occur. Otherwise, suppression will occur only if the suppression factor is greater than the fitness advantage of individuals at low population density. (**C**) The speed at which the TADS suppression drive reaches 99% of individuals in the population with varying introduction frequency and drive fitness. Full suppression or an equilibrium state will be attained within a few generations of this point. (**D**) Same as **C**, but with varying germline cleavage rate (the Y-linked drive can only be in males, so there would likely not be embryo cleavage - however, if there was paternal activity due to unusually high nuclease/gRNA expression or stability, this would be expected to further increase drive efficiency).

### Resistance to TA systems

With a modest degree of multiplexing, TA systems should generate substantially fewer resistance alleles than homing type drives without sacrificing drive performance, since there is no need for homology-directed repair. To study the rates at which r1 alleles (those which preserve the function of the target gene) are expected to form in such systems, we assumed that cleavage repair at a single site had a 10% probability of forming an r1 allele (instead of a disrupted allele), which appears to be near the upper end of the potential range of this parameter based on experiments^15–17^ (and by careful targeting, a significantly lower rate can be achieved^24^). The presence of a single disrupted site was considered to be sufficient to render the target gene disrupted, so to form a complete r1 allele, it was necessary to form an r1 sequence at each gRNA target site.

In drives with this high r1 formation rate, 100% efficiency, and assuming that drive homozygotes had a relative fitness of 95% compared to wild-type homozygotes, a single gRNA was not sufficient to allow for success of TARE, TADE, TADDE, or TADS same-site modification drives (Figure 9). Though all drives initially increased in frequency rapidly, the small fitness cost of the drive coupled with the high rate of r1 formation resulted in elimination of most drive alleles after 100 generations for TARE, TADE, and TADS. TADDE performed better, since r1 alleles would not be viable in the presence of a disrupted allele, while drive alleles would remain viable. Nonetheless, the final frequency of r1 alleles was still high for a scenario with only one gRNA. However, as the number of gRNAs increases, the number of r1 alleles that remain decreases drastically (Figure 9), indicating that for even very large populations, a modest number of gRNAs would likely be sufficient to preclude formation of resistance against the TA drives. Indeed, our calculations may substantially overestimate the number of r1 alleles formed, perhaps even greater than 100-fold. This is not only because we assumed a high proportion of repair resulting in r1 sequences, but also because the possibility for simultaneous cutting was not included in our deterministic model. However, such events should take place a substantial fraction of the time, particularly as the number gRNAs increases, and even one instance of simultaneous gRNA cleavage would likely cause a large enough deletion to prevent formation of an r1 allele. Additionally, homology-directed repair of drive cleavage using disrupted alleles as a template would likely preclude the formation of r1 alleles, which was also not taken into account in our model. Highly spaced gRNAs would reduce the chance of this taking place, but also increase the chance of successful disruption of the gene, making optimization of gRNA target spacing a potentially important consideration when designing these drives.

**Figure 9.**
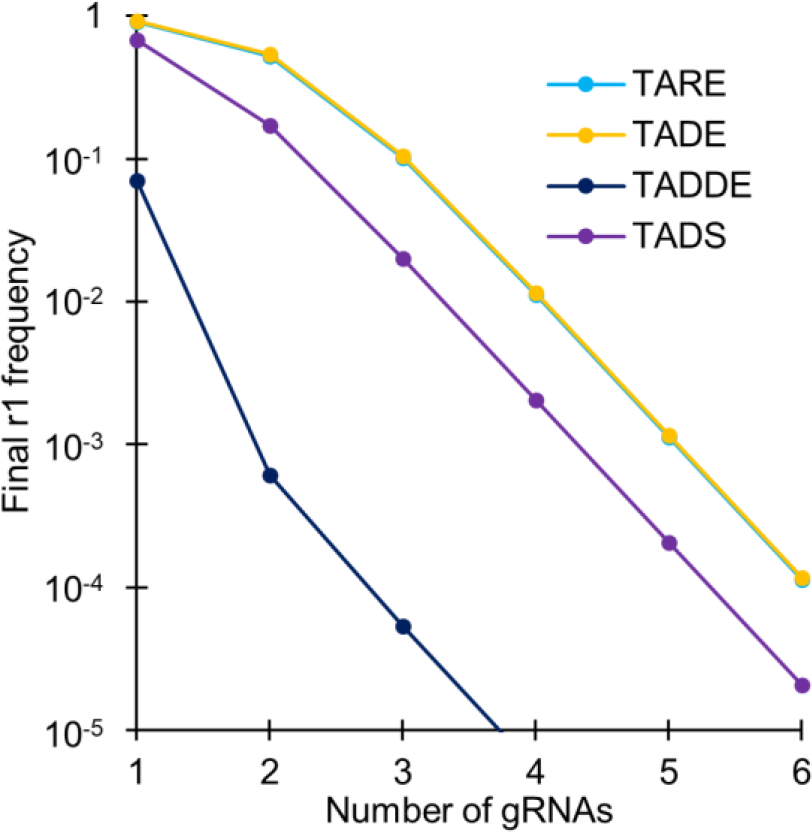
Resistance to TA systems. Simulations were conducted for drive systems with 100% cleavage rates (germline only for TADE) and 95% drive homozygote fitness. Each cleavage event was assumed to result in a functional r1 allele instead of a disrupted target allele with 10% probability. The number of gRNAs was varied, and a resistance allele was considered to be a functional “complete” r1 allele only if all gRNA cleavage sites possessed r1 sequences. The complete r1 allele frequency after 100 generations is reported.

## DISCUSSION

In this study, we have shown that TA gene drive systems should allow for the development of robust population modification or suppression drives that can be global or regionally confined. These systems have several major advantages over existing strategies (Table 1).

Most importantly, TA systems should be far less vulnerable to the formation of resistance alleles than homing drives. Multiplexing can somewhat ameliorate the formation of the more critical r1 alleles in homing drives, but this comes at the cost of reduced drive efficiency due to a variety of factors involved in homology-directed repair^16^. For population modification drives, it would likely be necessary for a homing drive to also target essential genes to reduce the rate of r1 resistance allele formation, opening up the possibility for partial homology-directed repair to continue to form r1 alleles, possibly at rates that would preclude success of the drive. TA drives are not expected to suffer from any efficiency loss upon multiplexing, allowing for effective elimination of r1 alleles given a sufficient number of gRNAs (Figure 9). Drive alleles are not typically copied in the early embryo by homology-directed repair, and only resistance alleles can form in this stage^15–17^. This gives homing drives a disadvantage compared to TARE, TADDE, and TADS drives, where cleavage events in the embryo would actually be beneficial for the drive. *Medea* avoids the formation of resistance by use of RNAi as the drive mechanism, and TA type drives could presumably be engineered similarly to use shRNAs or other RNAi taking effect during early embryo development, though this would somewhat limit the available array of potential gene targets to those with critical function at a developmental stage before the maternal RNAi is degraded.

Another potential advantage of TA systems, except TADS, is their threshold-dependent dynamics. This would have the effect of preventing establishment of the drive by occasional long-distance migration, thus confining the drives to a target region. While global drives may be desirable for reduction of vector-borne disease, regional confinement could often be important for political, economic, or conservation purposes^1,2,26^. TA systems allow for both regional population modification and suppression, giving scientists and policy-makers increased flexibility when considering the deployment of gene drives.

**Table 1.** Comparison of Drive Types.

| Drive Type | Speed | Threshold*/<br>Confinement | Promoter | Potential for<br>Resistance | Engineering<br>Difficulty |
| --- | --- | --- | --- | --- | --- |
| Homing | High | Zero | Germline | High | Low |
| Homing Suppression | High | Zero | Germline | Moderate## | Low |
| X-Shredder | High | Zero | Germline | Low | High |
| *Medea* | Medium | Low | Specific | Low | High |
| *Wolbachia*/2L2T** | Low | Medium | Any | Low | Low? |
| Underdominance | Low | High | Varies | Low | Low |
| TARE | Medium | Low | Any | Low | Low |
| TADE | Medium | Low | Germline | Low | Moderate |
| TADE Suppression | Medium | Medium | Germline | Low | Moderate |
| TADDE | Medium | Low | Any | Low | Moderate? |
| TADS | High# | Zero | Any | Low | High? |
| TADS Suppression | High# | Zero | Any | Low | High? |
| TADS Y Suppression | High# | Zero | Any | Low | High? |

\*Thresholds assume a small fitness cost for the drive. Threshold is an indirect measure for the degree of confinement. Zero-threshold drives will potentially spread globally, low and medium levels of confinement will constitute regional drives (possibly local for larger fitness costs), and high threshold systems should remain in a local area, if they are able to successfully persist<sup>34</sup>.
\*\*A 2-Locus 2-Toxin-antitoxin (2L2T) underdominance design with two TARE-like alleles.
#The speed of a TADS drive is reduced if target species females can mate multiple times and sperm from different males compete to fertilize eggs.

Aside from the configurations presented in this manuscript, the same principles could potentially be applied to other designs. For example, to achieve a greater degree of local confinement (at the costs of greater release sizes, as is usually the case with such systems), a 2-locus 2-toxin-antidote system could be engineered by using two TARE systems, each of which providing rescue for the target gene of the other system. Such a system could presumably be engineered quite easily and combined with a tethered^33^ homing suppression drive. Note, however, that the germline-only nuclease promoter needed for homing in this case may slow down the TA underdominance drive due to lack of embryo activity. An alternative would be to use two different nucleases in the underdominance TARE components. Local suppression could also be obtained with a similar 2-locus TADE system, with one of the TADE alleles disrupting a sex-specific fertility gene, as in TADE suppression. Indeed, a standard TADE suppression system with a promoter that has high embryo activity could itself be a feasible method for local population suppression. Alternatively, either of these TADE methods could be used for modification if not located in a fertility gene. Indeed, a TARE drive with a target that is not fully haplosufficient might also constitute an effective underdominance-type drive. Each of these systems should be possible to engineer with current techniques and targets that are already available.

TA systems have a high degree of flexibility regarding target genes. TARE systems would likely be most efficient when using recessive lethal targets that take effect in the early embryo, to reduce competition for surviving drive-bearing offspring. However, other recessive lethal or sterile targets (including sex-specific) could also be used. TADE targets should be haplolethal, but a high degree of haploinsufficieny will usually also be tolerable, thus allowing for some flexibility in target selection. On the other hand, TADS targets are highly specific, and this will likely be the limiting factor in engineering these systems. Several genes have been found in *Drosophila melanogaster* with post-meiosis I expression in males^35–37^, yet it remains to be seen if a gene can be identified for which expression in this interval is necessary for successful completion of spermatogenesis, thus making it a potential TADS target. It should also be noted that while efforts to design a successful X-shredder system have been stymied by low transgene expression from the Y chromosome, TADS Y-suppression systems may not suffer from this, since they would only need to cleave a few targets in a single gene, rather than dozens of targets simultaneously across an entire chromosome.

Overall, our study shows that TA systems can provide flexible and effective mechanisms for a variety of potential gene drive applications. Their feasibility has already been demonstrated for TARE in *D. melanogaster* for same-site^29^ and distant-site^30^ configurations. Future experiments should investigate the feasibility and dynamics of the other TA drives we have proposed here and how they can be brought to other species of interest such as mosquitoes. Modeling should further explore how these proposed systems would behave in more realistic, spatially structured populations.

## ACKNOWLEDGEMENTS

This study was supported by the National Institutes of Health award R01GM127418 to P.W.M., National Institutes of Health award R21AI130635 to J.C., A.G.C., and P.W.M, and the National Institutes of Health award F32AI138476 to J.C.

## SUPPLEMENTAL INFORMATION

**Figure S1.**
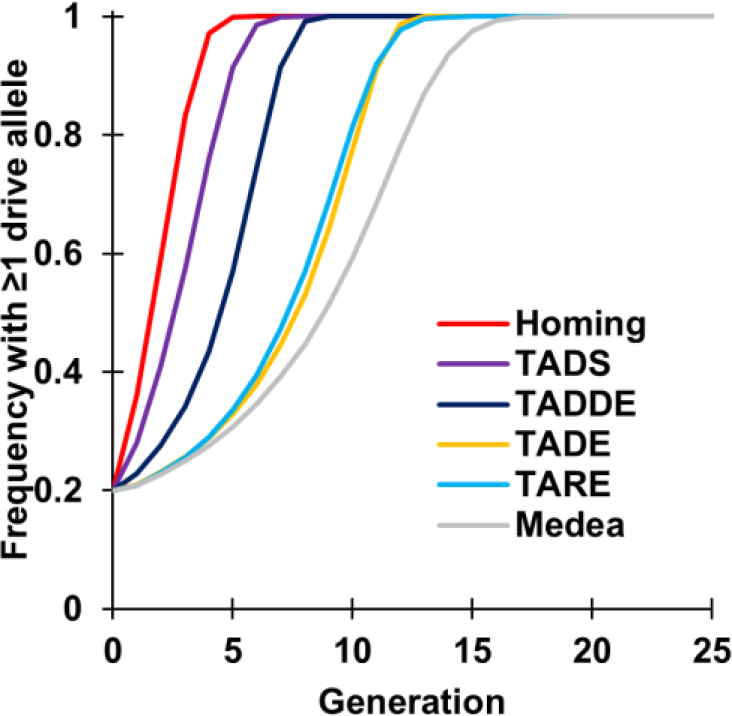
Frequency of individuals with at least one drive allele. Drive heterozygote individuals are released into a wild-type population at 20% starting frequency (10% starting drive allele frequency) in a discrete-generation, deterministic model. All drives are assumed to have no fitness cost and 100% drive efficiency. The frequency of individuals with at least one drive allele is shown.

**Figure S2.**
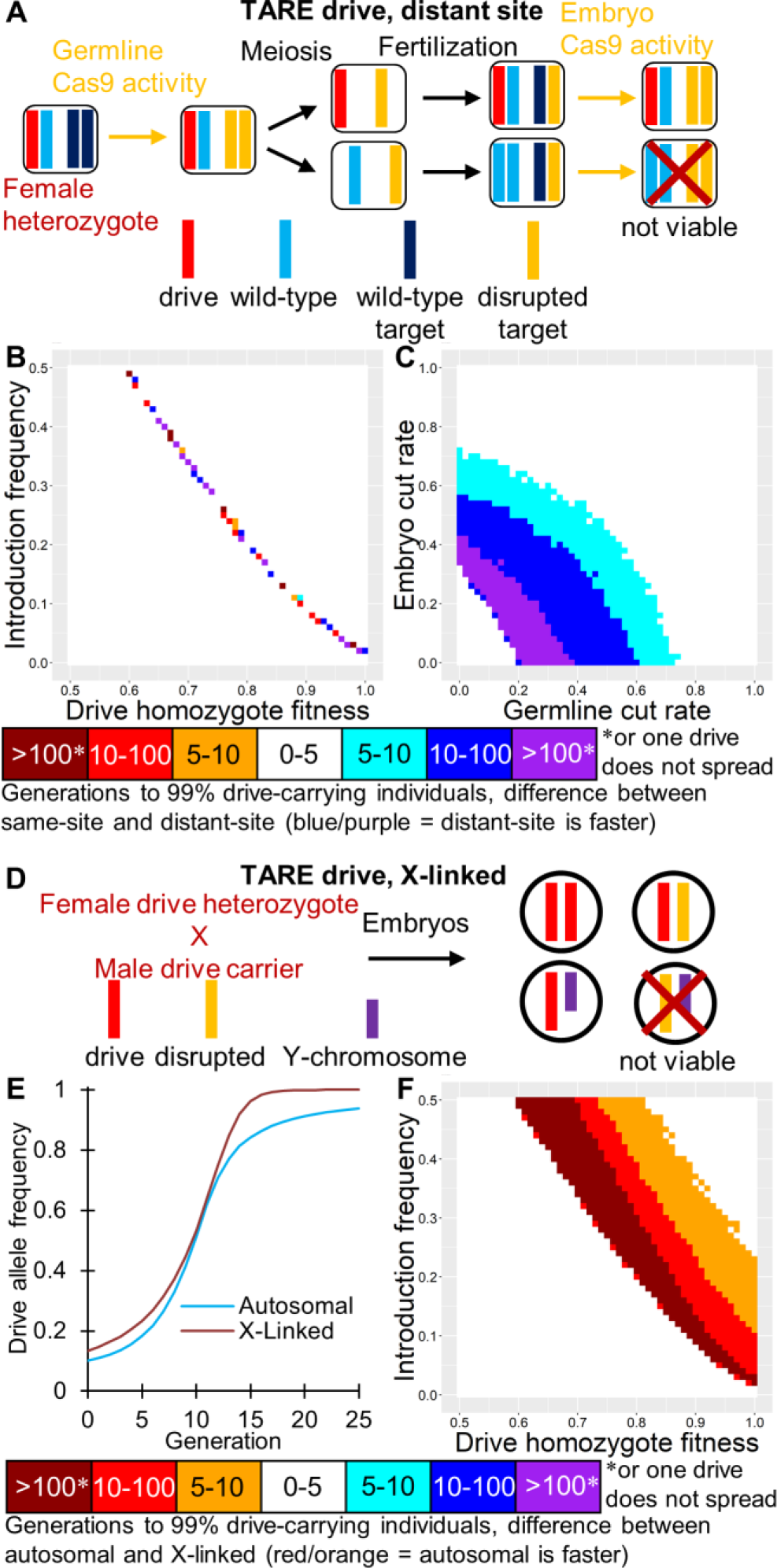
Distant-site and X-linked TARE. (**A**) In the distant-site TARE drive, the target gene is unlinked from the drive allele. Individuals with two disrupted target alleles without any drive alleles are not viable. (**B**) The speed at which the distant-site TARE drive reaches 99% of individuals in the population with varying introduction frequency and drive fitness compared to the same-site drive. (**C**) Same as **B**, but with varying germline and embryo cleavage rate. (**D**) In an X-linked TARE drive, males with only one disrupted copy of the target gene are not viable. (**E**) This allows an X-linked drive to reach fixation substantially more quickly than and autosomal TARE drive. (**F**) The speed at which the same-site X-linked TARE drive reaches 99% of individuals in the population with varying introduction frequency and drive fitness compared to the autosomal same-site drive.

**Figure S3.**
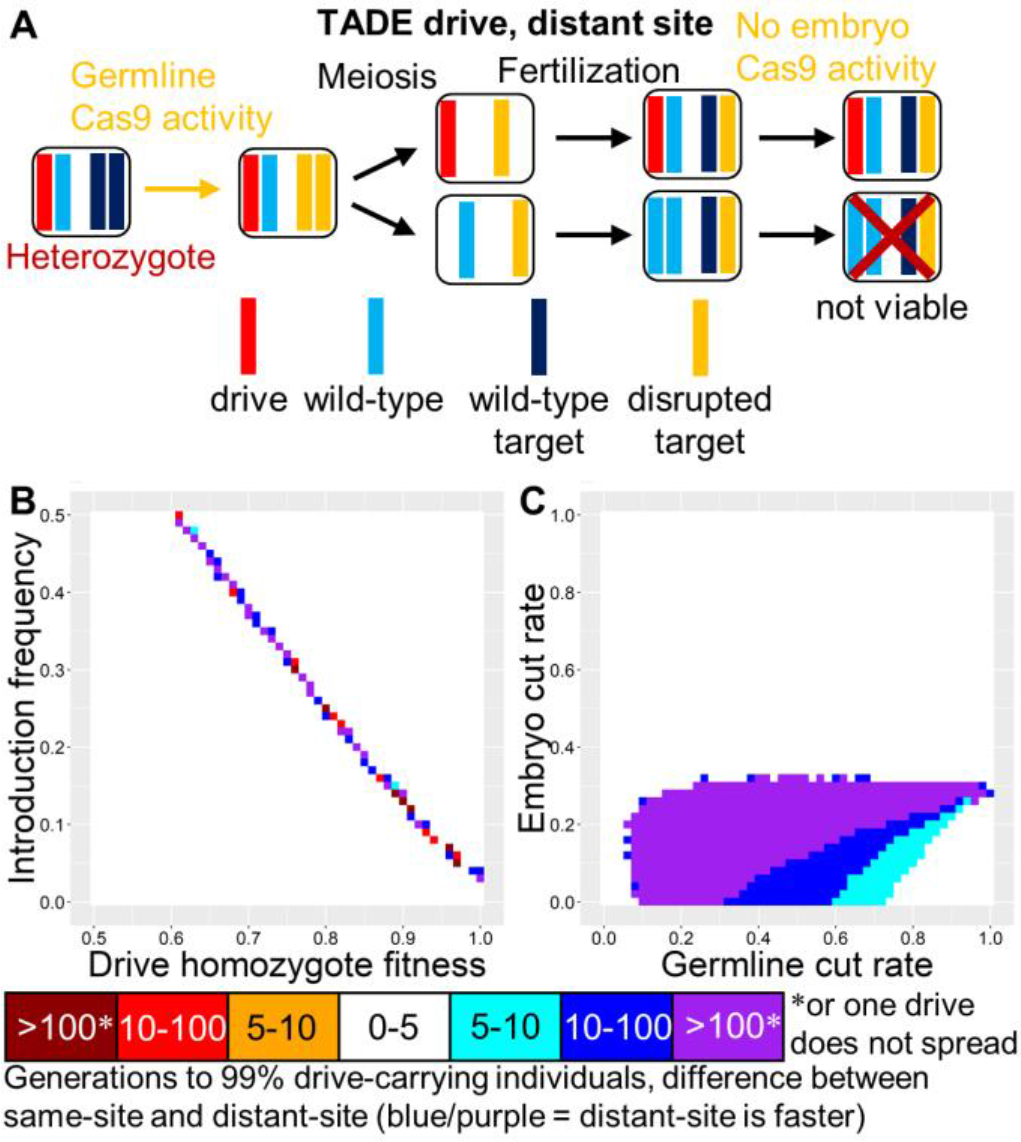
Distant-site TADE. (**A**) In the distant-site TADE drive, the target gene is unlinked from the drive allele. Individuals with fewer than two wild-type target alleles plus drive alleles not viable. (**B**) The speed at which the distant-site TADE drive reaches 99% of individuals in the population with varying introduction frequency and drive fitness compared to the same-site drive. (**C**) Same as **B**, but with varying germline and embryo cleavage rate.

**Figure S4.**
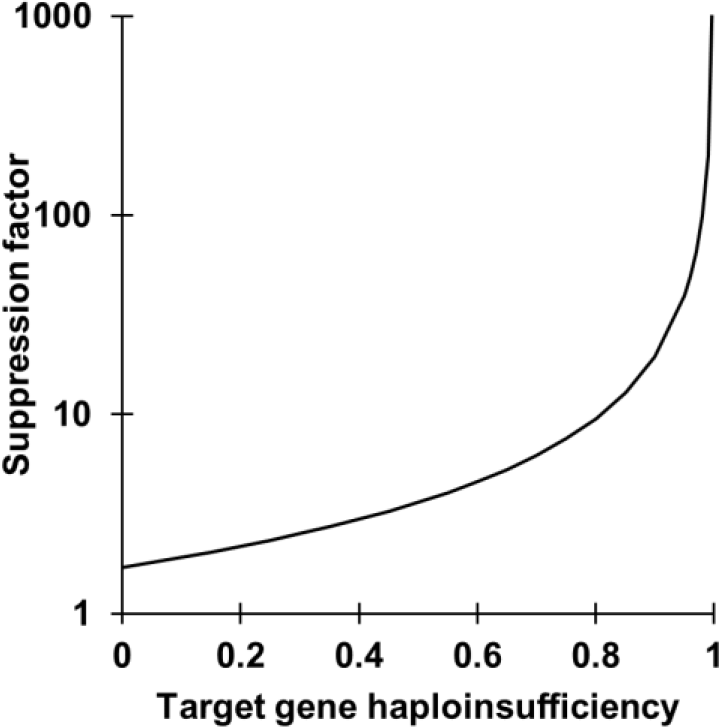
Haploinsufficiency in TADE suppression. If the target allele is not haplolethal but merely haploinsufficient, a TADE suppression drive will impose a suppression factor on the population, which may not cause complete suppression, depending on species and ecological parameters, as well as the degree of haoloinsufficiency.

**Figure S5.**
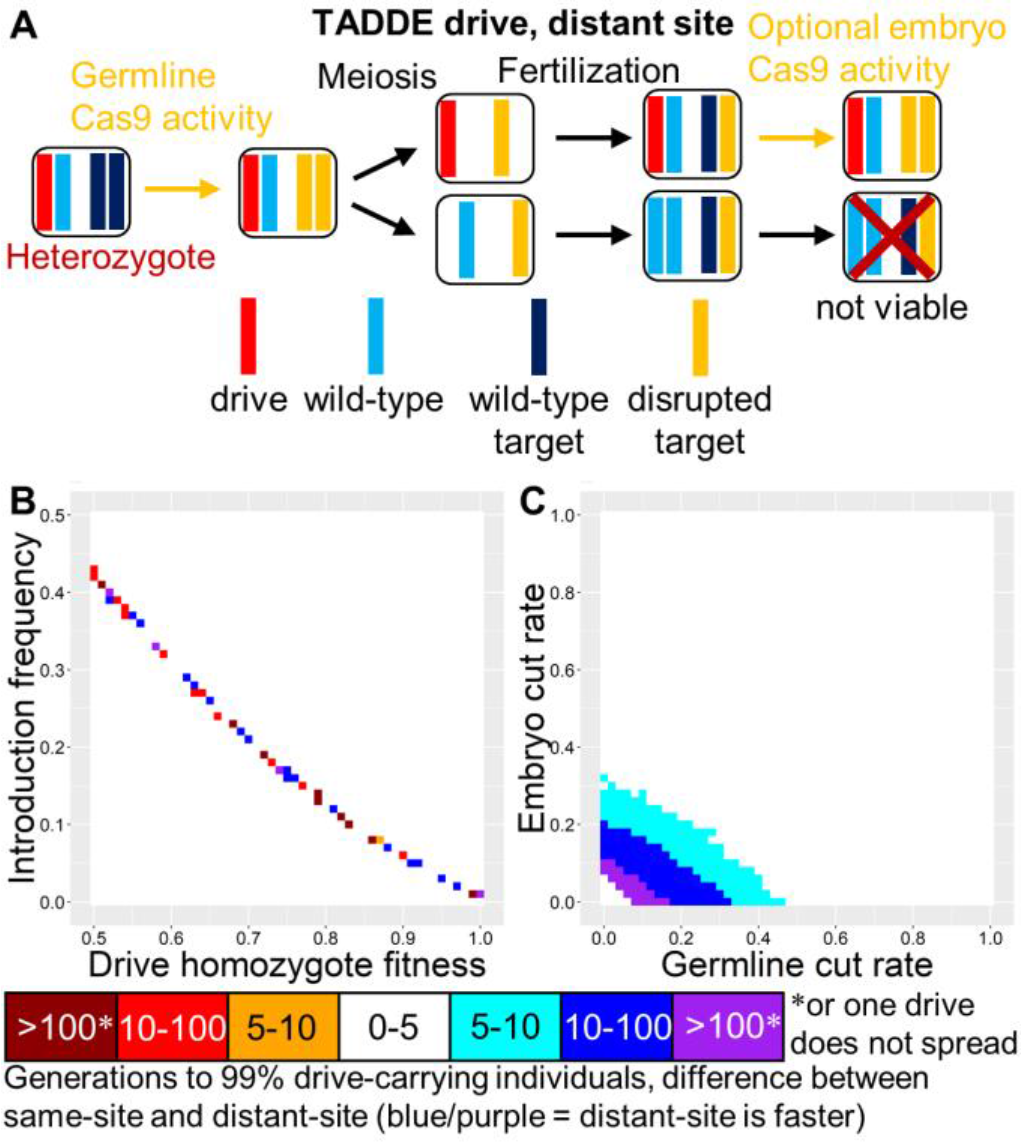
Distant-site TADDE. (**A**) In the distant-site TADDE drive, the target gene is unlinked from the drive allele. Individuals with at least one disrupted target allele are not viable unless they have at least one drive allele. (**B**) The speed at which the distant-site TADDE drive reaches 99% of individuals in the population with varying introduction frequency and drive fitness compared to the same-site drive. (**C**) Same as **B**, but with varying germline and embryo cleavage rate.

**Figure S6.**
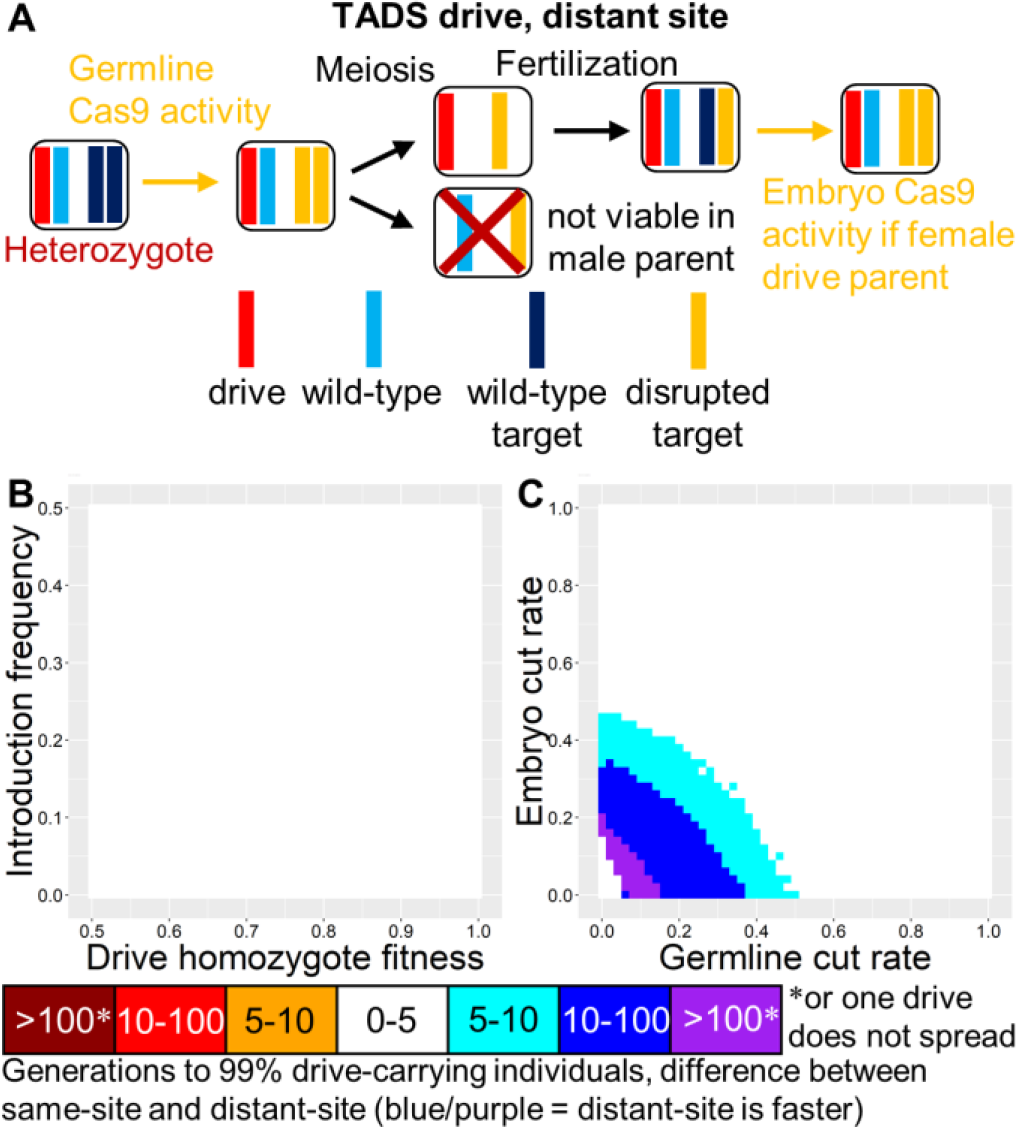
Distant-site TADS. (**A**) In the distant-site TADS drive, the target gene is unlinked from the drive allele. Sperm with a disrupted target allele are not viable unless they also have a drive allele. (**B**) The speed at which the distant-site TADS drive reaches 99% of individuals in the population with varying introduction frequency and drive fitness compared to the same-site drive. (**C**) Same as **B**, but with varying germline and embryo cleavage rate.

